# Integrating various Experimental Information to Assist Protein Complex Structure Prediction by GRASP

**DOI:** 10.1101/2024.09.16.613256

**Authors:** Yuhao Xie, Chengwei Zhang, Shimian Li, Xinyu Du, Min Wang, Yingtong Hu, Sirui Liu, Yi Qin Gao

## Abstract

Protein complex structures are essential for understanding of biological activities and drug development. Improving complex structure prediction accuracy of AI models for cases such as antigen-antibody complexes is expected to further enhance their applicability. Meanwhile, a large variety of experimental methods are used to provide structural insights for protein complexes, with only sparse or approximate knowledge obtained. A general tool is needed to integrate AI models with limited experimental information for high-throughput and accurate protein complex structure prediction. To efficiently and flexibly incorporate the different forms of experimental information, we introduce here GRASP. GRASP outperforms existing tools in handling both simulated and real-world experimental restraints including those obtained from XL, CL, CSP, and DMS. As an example, GRASP excels in predicting antigen-antibody complex structures, even surpassing AF3 when utilizing experimental DMS and CL restraints. In addition to accelerating the restrained modeling process, its ability to integrate multiple forms of restraints makes it capable of integrative modeling. We also showcase its potential in modeling protein structural interactome in the near-cellular condition based on large-scale *in vivo* XL data for mitchondria.

## Introduction

Protein complexes are essential in multiple biological processes^1^. The disparity between known amino acid sequences and solved protein structures highlights the critical need for computational protein structure prediction^2,3^. However, current structure prediction approaches such as AlphaFold-Multimer(AFM)^4^, ESMFold^5^, and OmegaFold^6^ still show limited accuracy in protein complex structure prediction^7^.

Meanwhile, various experimental approaches exsit to provide sparse information to assist predict protein complex structures. Residue pair restraints (RPRs) can be obtained by methods such as cross-linking mass spectrometry (XL-MS)^8,9^ and Nuclear Magnetic Resonance Nuclear Overhauser Effect Spectroscopy (NMR NOESY)^10^, which provide distance restraints between residue pairs. Interface restraints (IRs) are used to identify interface residues of the complex. Differential covalent labeling (CL) detects reduced covalent modifications when protein complexes transition from unbound to bound states^11^. Chemical shift perturbation (CSP) identifies regions interacting with ligands by observing shifts in chemical signals upon binding^12,13^. Deep mutational scanning (DMS) highlights mutations affecting binding, often corresponding to interface residues^14,15^. These methods all provide information on IRs. In addition, *in situ* biophysical experiments including cryo-ET^16^, solution NMR^17^, *in vivo* XL^18^ and other techniques develop rapidly to detect protein interactions at their real functional state. Integrating these experimental data to structure modeling will tremendously benefit the understanding of complicated biogical systems.

Several tools were developed to integrate experimental data into protein structure prediction. For instance, AlphaLink^19^ embeds crosslinking information directly into the pair representation before it enters the Evoformer block and has a protein complexes version^20,21^. ColabDock^22^ leverages experimental restraints for protein-protein docking by inverting AlphaFold models but is relatively time-consuming and is limited to proteins under 1200 residues. Drake et al^23^ integrates a covalent labeling scoring term into RosettaDock score function to select models aligned with experimental data, but is limited to CL data. Besides deep learning based approaches, HADDOCK^24,25^ and ClusPro^26–28^ are two representative traditional protein-protein docking methods which can integrate experiment restraints in different ways. HADDOCK transforms the restraints into energy term to guide conformation sampling and scoring while ClusPro uses restraints information to filter out FFT-generated structure pool.

Additionally, since different experimental methods provide cross-validated, or complementary information, they can be notably helpful for modeling of complicated complexes. People have developed integrative modeling approaches to handle this, a representative approach is the Integrative Modeling Platform (IMP)^29^ which samples conformation protein complex through Monte Carlo simulation and estimates the conditional likehood of structures given restraints in the bayesian framework. For now the integrated modeling approaches are mostly case specific and need much manual effort^30,31^.

To address these challenges, we introduce here a protein complex structure prediction model called Generalized Restraints Assisted Structure Predictor (GRASP). GRASP integrates both RPR and IR restraints directly into the model to accommodate diverse experimental data in model inference. It is fine-tuned with restraint-related loss terms and applies iterative noise filtering in inference to reduce the impact of erroneous restraints. We perform comprehensive evaluations on both simulated and real-world experimental data on varied functional protein complexes to assess GRASP’s ability to handle various types of experimental information, both simultaneously and independently, to improve the accuracy of protein complex structure predictions. Specifically, we show the application of GRASP to antibody-antigen complex structure determination, integrative modeling that simultaneously makes use of multiple experiemtnal data, and prediction of *in situ* complex structures captured in mitochondria. In this way, GRASP can help provide reliable information on structures and interactions of complex biological machines, potentially also molecular details on dynamical protein interaction in cellular environment when information such as XL data is provided.

## Results

### GRASP integrates two types of general restraints into protein structure prediction model

Both RPRs and IRs are integrated into AlphaFold-multimer (AFM) for protein complex structure prediction by GRASP. RPRs, derived from experimental techniques like XL-MS and NMR, impose distance restraints between residue pairs and can be viewed as edge features in a protein complex graph with residues treated as nodes. IRs, obtained from experimental methods like CL, CSP, and DMS, are integrated into the model as node features, indicating whether a residue is at the interface based on experimental evidence. These features are incorporated into AFM via three blocks (Fig. 1a). The RPRs are integrated into GRASP as MSA bias and IPA bias (Fig. 1b) following our previous work^32^. To incorporate IRs into our network, we use relative position encoding and introduce interface fusion blocks within the Evoformer and IPA modules (Fig.1c).

**Fig. 1.**
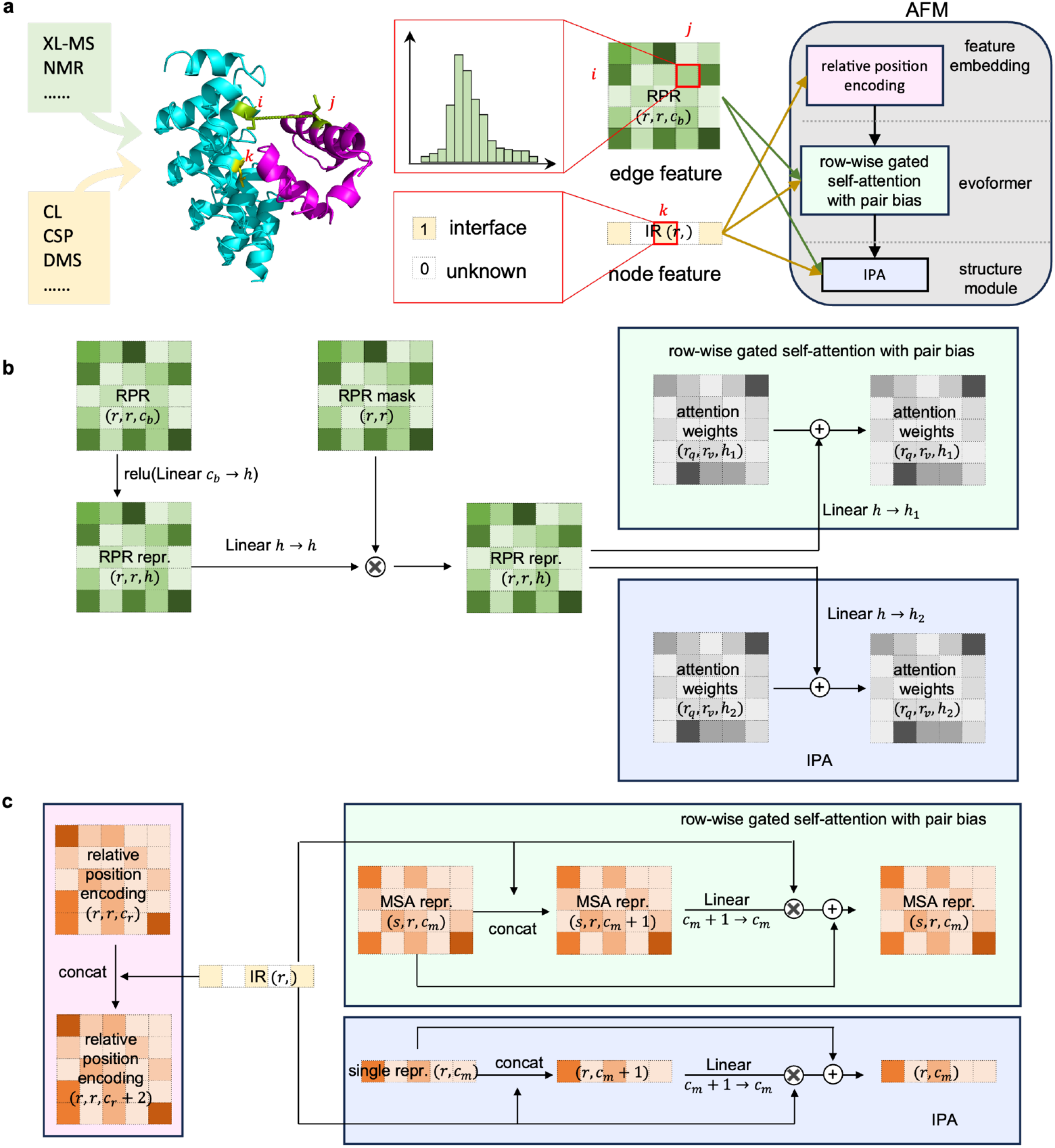
Scheme of GRASP to integrate generalized restraints. **a**, The GRASP architecture shows how two types of restraints from various experiments are integrated using three AFM blocks. **b-c**, Details of RPR (**b**) and IR (**c**) integration are illustrated, with arrows indicating information flow and array shapes noted in parentheses.

We added four restraint-related loss terms (see Methods) to the original loss function and fine-tuned GRASP for 22,000 steps using 64 Ascend 910A processors starting from the AFM v2.3 model-1 checkpoint. Four checkpoints are selected in the finetuning process for inference. In addition, to provide more diversified checkpoints, we trained a multimer prediction model from scratch to reach a baseline complex prediction accuracy comparable to AFM (see Methods and Fig. S1), and applied the same fine-tuning strategy for 22,000 steps to obtain the fifth model checkpoint. In standard GRASP inference, similar to AFM, each of the five models was run with five random seeds, with the best predictions selected based on pLDDT scores and restraint recall. We also applied an iterative noise filtering strategy to reduce the impact of erroneous restraints (see Methods and Fig. S2).

### GRASP effectively integrates RPRs and IRs on self-curated test dataset

To evaluate GRASP’s performance with the two independent types of restraints, we curated a benchmark dataset by selecting samples that were released after training set samples. The benchmark set differs from the training set in chain composition, have no internal redundancies, and show suboptimal AFM predictions (see Methods), to prevent data leakage and ensure dataset diversity as well as quality. The curated dataset contains 713 interfaces from 313 challenging complexes. For RPRs, simulated restraints were sampled from inter-chain residue pairs within 8 Å (contact-RPR). For IRs, restraints were sampled from interface residues which are within 8 Å of the other chain in the dimer. IRs are less informative than RPRs as they only provide potential contact residues from one protein. We incorporated varying numbers of simulated restraints and compared the performance of AlphaLink, ClusPro, HADDOCK, ColabDock and GRASP on their applicable restraint types.

As the number of restraints increased, the prediction accuracy of the restraint-based methods improved consistently for both types of restraints. GRASP consistently ranked first among all methods tested and delivered the most substantial gains across different quantities of both IRs and contact-RPRs (Figs. 2a-b, Fig. S3). With just two inter-chain contact RPRs, GRASP reached a mean DockQ of 0.35, with 52.7% of samples exceeding the acceptable threshold of 0.23, demonstrating its effectiveness even with minimal restraints. For the less informative IRs, GRASP achieved mean DockQ scores and success rates of 0.24 and 35.3%, 0.34 and 51.9%, and 0.41 and 63.2% for 4, 10, and 20 restraints, respectively.(Fig. 2a-b, Table S1). AlphaLink showed similar performance to AFM with contact-RPRs, indicating its limitation in handling different crosslinking distances. For both RPR and IR, ColabDock, HADDOCK and ClusPro showed overall better performance than AFM, with ColabDock performing slightly better among the three.

**Fig. 2.**
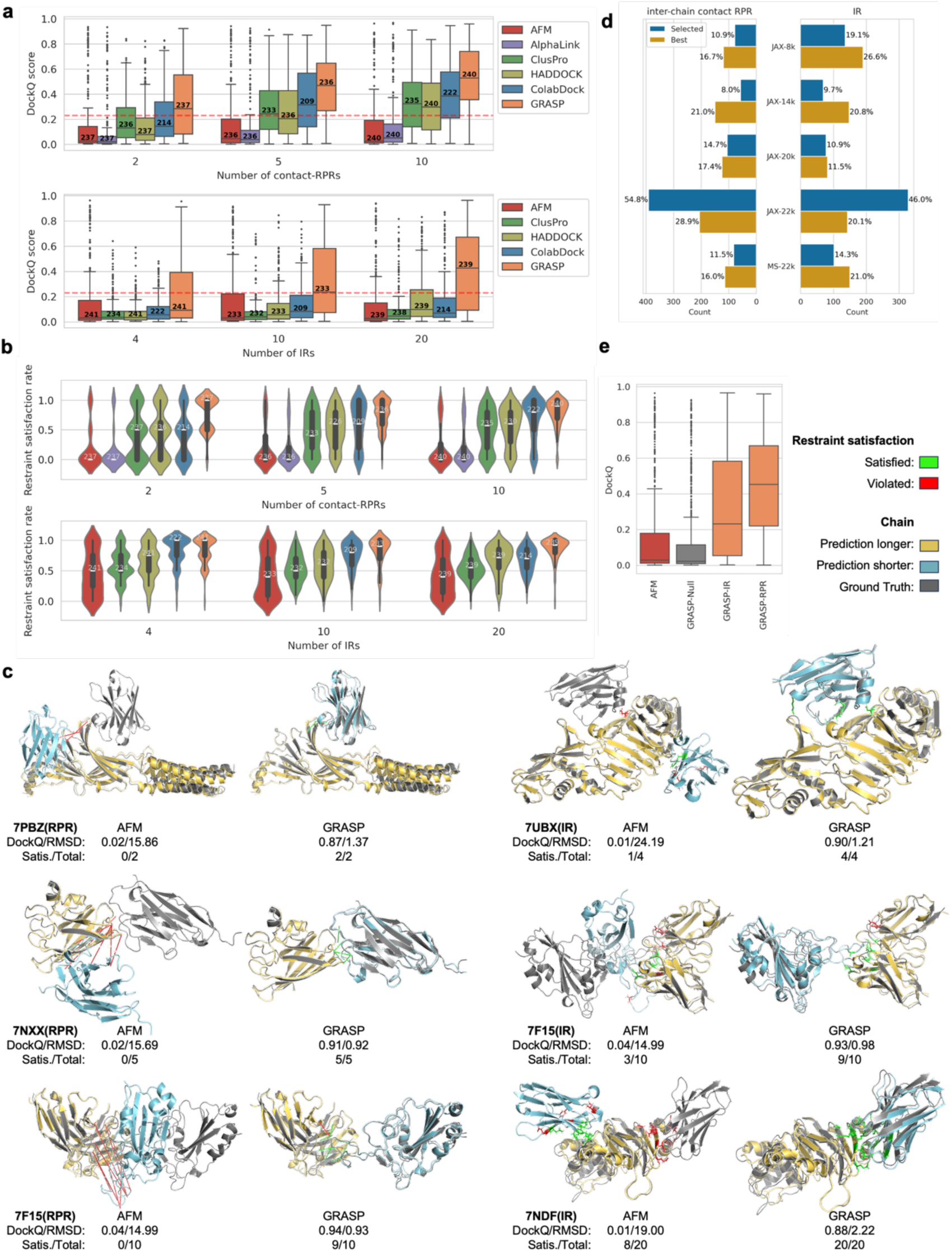
Performance of different methods on benchmark dataset with two types of basic restraints. **a-b.** Comparison of DockQ (**a**) and restraint satisfaction rate (**b**) between different methods for varying numbers of inter-chain contact residue pair restraints (upper) and interface restraints (lower). The red dashed horizontal lines in each subfigure indicate a DockQ value of 0.23.The numbers above the median line in the boxplots and violin plots represent sample sizes. ClusPro fails in some samples, and ColabDock couldn’t predict proteins with more than 1200 residues, leading to smaller sample sizes compared to others. **c**.Representative structures of GRASP and AFM on selected samples. For each structure DockQ/RMSD and the number of satisfied restraints/total restraints are shown in the figure. All short chains in the predicted structures are colored cryo, while long chains are yellow, superimposed on the real structure (gray). Green indicates satisfied restraints, while red shows violated ones. All IRs are displayed as sticks, while restrainted residue pairs are connected by solid lines. AFM was used without any restraints, GRASP restraints are also shown on the AFM structure only for ease of comparison. **d**. The proportion of the selected structures and the best structures according to DockQ from the five models in GRASP. **e**. Ablation study for GRASP performance with and without restraints. GRASP without restraints (GRASP-Null) showed no improvement and is comparable to AFM. In all boxplots, the central lines indicate the median, the box edges show the quartiles, and the whiskers extend to 1.5 times the interquartile range, with points beyond this being outliers.

The protein complex of PDB ID 7PBZ is a representative case. The prediction of AFM yielded a DockQ of 0.02. With only two contact-RPRs, GRASP was able to find the correct interface and improved the DockQ to 0.87. Similarly, in the case of 7UBZ, GRASP raised DockQ score from the AFM DockQ of 0.01 to 0.9 with four IRs. In other examples including 7NXX, 7F15, and 7NDF, significant improvements are also observed with GRASP incorporating simulated restraints (Fig. 2c).

We further performed ablation analysis to evaluate the contribution of models and restraints. To evaluate the contribution of the five models, we examined the proportion of final selections and best predictions across the five models. We found that each model contributed to the final selections, with the JAX-22k checkpoint contributing the most. Compared with the selected structures, the best structures were more evenly distributed among the models (Fig. 2d). Similar to observations in AFM, using multiple models and selecting the top-scoring prediction enhanced both accuracy and robustness, and reduced the risk of failure from a single model (Fig. S4).

We also evaluated the model performance without restraints. GRASP shows comparable accuracy to AFM, with mean DockQs of 0.15 and 0.17, respectively (Fig. 2e). This result indicates that the improvements by GRASP stem from its ability to incorporate restraints rather than just fine-tuning. Additionally, the chain length showed a weak negative correlation with prediction accuracy, which is expected since larger proteins require more restraints for accurate predictions. (Fig. S5).

### GRASP improves structure prediction assisted by diverse single restraint types: XL, CL, and CSP

In this section, we benchmark the performance of GRASP and other methods on XL, CL, or CSP data,.

Chemical crosslinking mass spectrometry (XL-MS) data is one of the most common experiments to obtain residue pair restraint information. To evaluate the model ability of integrating XL-MS data, we simulated two sets of SDA cross-linking restraints on the 294 complexes from the self-curated dataset and collected experimental XL-MS data for protein complexes. In the first set, the number of restraints was set to 2% of the total sequence length, with this restraint coverage based on the work of Bartolec et al^33^. In the second set, this ratio was increased to 5% to simulate mixture of several crosslinking reagents. In both sets, the maximum restraint distance was 25 Å, with 20% of the restraints violating the threshold to mimic real-world experimental XL-MS data (see Methods). Since ColabDock can only handle restraints below 22 Å, and ClusPro can only allow restraints between two rigid chains or subcomplexes, we evaluated GRASP, AlphaLink, and HADDOCK on these two simulated datasets.

Overall, GRASP demonstrated superior performance for both simulated XL restraint conditions, consistently outperforming HADDOCK and AlphaLink in both accuracy and restraint satisfaction (Fig. 3a, Table S2). For comparison, AFM achieved a mean DockQ score of 0.07. Under the 2% restraint condition, GRASP exhibited an improvement in prediction accuracy, with a mean DockQ score of 0.21, outperforming both HADDOCK (0.08) and AlphaLink (0.11). GRASP also achieved higher restraint satisfaction and correct restraint satisfaction rates, with medians of 69.2% and 81.3%, respectively, compared to AlphaLink’s 42.3% and 50.0%, and HADDOCK’s 61.1% and 71.4%. Although HADDOCK’s performance in DockQ was inferior to AlphaLink, its structures better satisfied the input restraints. Even with 20% erroneous restraints, GRASP achieved over 80% satisfaction of correct restraints, indicating its robustness against experimental noise. When the restraint coverage increases to 5%, GRASP, HADDOCK and AlphaLink all improve to reach mean DockQs of 0.27, 0.10, and 0.17, respectively, and GRASP still achieves the highest accuracy. The single-model results of GRASP already surpassed AlphaLink, and further improvements are observed when multiple models are used (Fig. S6). Interestingly, although AlphaLink was fine-tuned on simulated SDA restraints^20^, it did not perform as well as the more general-purpose GRASP, suggesting that GRASP may have learned robustly the physical features.

**Fig. 3.**
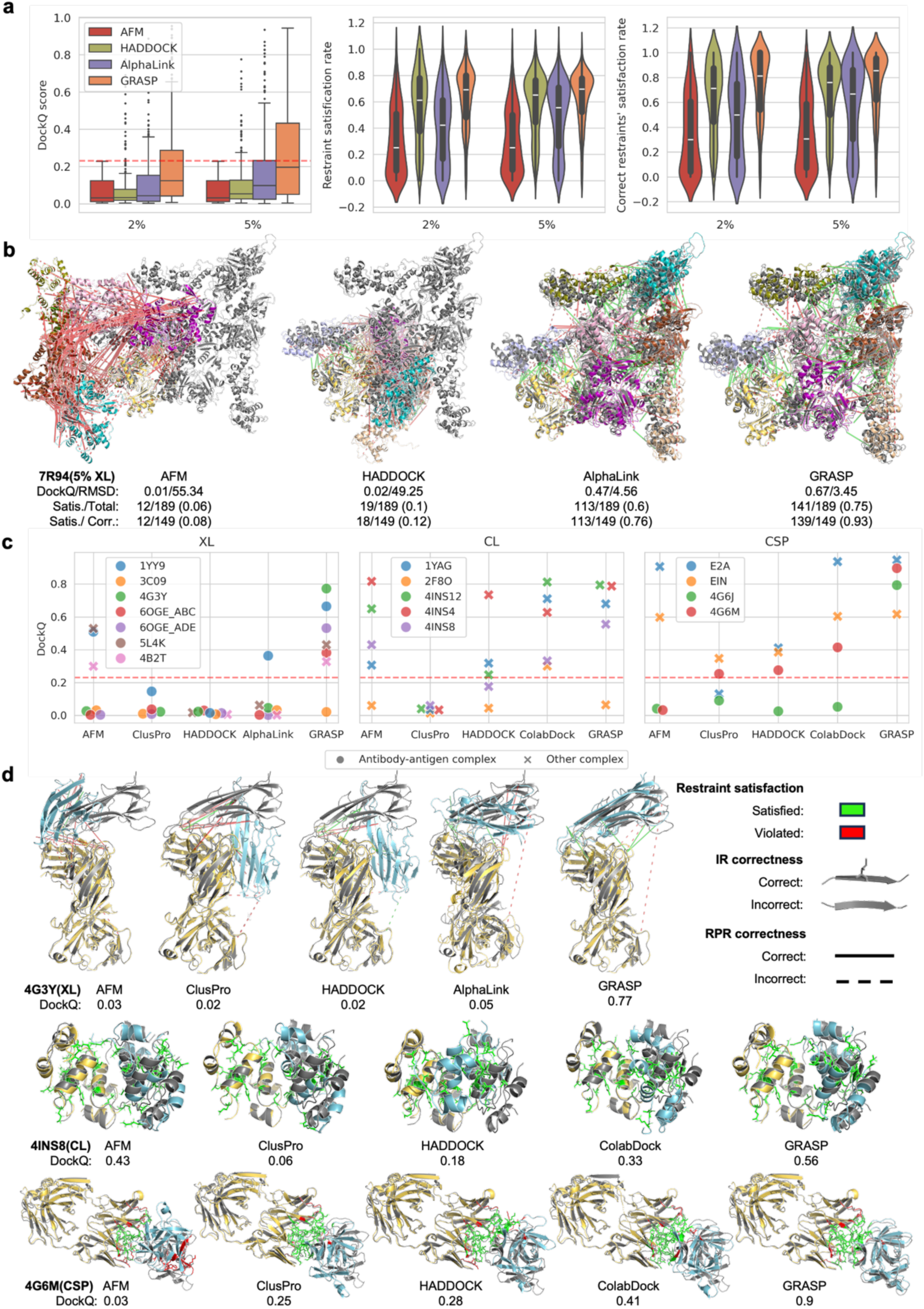
The performance of GRASP and other methods using single-type experimental restraint data. **a.** DockQ (left), restraint satisfaction rate (middle), and correct restraint satisfaction rate (right) for various methods on simulated SDA cross-linking mass spectrometry data across different restraint coverages. AFM did not use restraint data. In the boxplot, the central lines indicate the median, the box edges show the quartiles, and the whiskers extend to 1.5 times the interquartile range, with points beyond this being outliers. The red dashed horizontal line in the left subfigure indicates a DockQ value of 0.23 (Same for subfigures in panel **c**). **b.** 3D visualization of predicted structures by various methods compared to the true structure (gray) on 7R94 in the simulated XL dataset. **c.** DockQ scores for various methods using single-type real-world experimental restraint data, including XL (left), CL (middle), and CSP (right). Since the number of cases for each experimental approach is lower than 10, all of the evaluated cases are drawn separately as scatters for different methods. **d.** 3D cartoon of representative predicted and true structures for each method under different experimental restraints. The predicted structures are colored cyan (smaller part) and yellow (larger part), with the latter superimposed on the real structure in gray. Satisfied restraints are shown in green and violated ones in red. Correct IRs are displayed as sticks, incorrect ones as cartoons. Correct RPRs are connected by solid lines, while incorrect RPRs are shown with dashed lines. AFM was used without restraints, while GRASP restraints are displayed on the AFM structure for comparison.

GRASP can also perform well on large complexes, such as 7R94, which has over 3,500 residues (Fig. 3b). In this case, AFM mispositioned many chains, and even after adding simulated XL restraints, HADDOCK could not rescue the prediction. Both AlphaLink and, especially, GRASP showed significant improvements, yielding DockQ scores of 0.47 and 0.67, respectively.

We gathered real-world experimental restraint data from various studies (see Methods and Table S3) and tested how different methods leverage these restraints on the corresponding datasets. We collected seven samples with XL data^34–37^ (Fig. 3c left). GRASP achieved a mean DockQ score of 0.45, outperforming other tools such as AFM (0.20), ClusPro (0.04, based on five samples), HADDOCK (0.02), and AlphaLink (0.07). For the two non-antigen-antibody complexes (4B2T and 5L4K), both GRASP and AFM made successful predictions with DockQ higher than 0.23, while AlphaLink and HADDOCK failed. Among the five antibody-antigen XL examples, AFM had only one prediction with DockQ score above 0.23 but with an RMSD of 26.37, indicating that only the interface region was aligned closely with the true structure (Fig. S7). After providing experimental XL data, GRASP showed significant improvements in 80% of the cases. In the case of 4G3Y, with 12 DSBU (30 Å cross-link distance) restraints including three erroneous ones, GRASP increased the DockQ score to 0.77 from a value of 0.03 for AFM, satisfying all nine correct restraints while ignoring the three incorrect ones. In contrast, AlphaLink satisfies only two restraints while HADDOCK was misled by the incorrect restraints, both resulting in a DockQ score below 0.23. This observation indicates that GRASP not only ensures structure alignment with the provided restraints, but also excel in restraint discrimination abilities, minimizing the impact of erroneous restraints.

For five cases with experimental CL data from Drake et al^23^, AFM achieved an average DockQ of 0.45, while HADDOCK and ClusPro show average DockQ values of 0.3 and 0.04, respectively. ColabDock improved to an average DockQ of 0.56, and GRASP showed the greatest improvement, reaching an average DockQ of 0.58 (Fig. 3c middle). Notably, for 4INS8, GRASP (DockQ of 0.56) is the only method that outperforms AFM (DockQ of 0.43) (Fig. 3d). We gathered four CSP cases from two studies^25,38^ (Fig. 3c right). AFM performed well in two cases but struggled with the remaining two antibody-antigen complexes, resulting in an average DockQ of 0.39. HADDOCK and ClusPro showed average DockQ values of 0.27 and 0.21, respectively. ColabDock improved the performance to 0.5, while GRASP further raised the average DockQ to 0.81. Notably, GRASP accurately predicted 4G6J with a DockQ of 0.79, whereas all other methods failed. In another challenging case, 4G6M, GRASP improved the prediction to 0.9 using CSP data. ClusPro, HADDOCK, and ColabDock also improved but only reached DockQ scores of 0.25, 0.28, and 0.41, respectively (Fig. 3d). It is worth noting that all methods except for GRASP performed poorly on these antibody-antigen complexes, highlighting the challenge of this task and the need for focused efforts in this area (Fig. 3c-d).

### GRASP performance on Antibody-antigen prediction

Structure prediction of antigen-antibody complexes is critical for antibody design and immunotherapy. However, current methods like AFM have limitations in accuracy. AF3^39^ has been reported to perform better on antigen-antibody complexes, and we included it in our evaluation in this section. Since it only supports manual server submission process with daily task limits, we assessed AF3 only on collected samples with real-world experimental restraints.

We first evaluated GRASP on the BM5.5^40^ antibody-antigen dataset. This dataset comprises 67 samples, each equipped with three sets of simulated DMS interface restraints (see Methods). Results in Fig. 4a and S8 (also Table S4) showed that AFM outperformed ClusPro, HADDOCK, and ColabDock, achieving a DockQ median of 0.25 and a success rate of 52.2%. GRASP, however, surpassed all these methods with a DockQ median of 0.64 and a success rate of 71.6%.

**Fig. 4.**
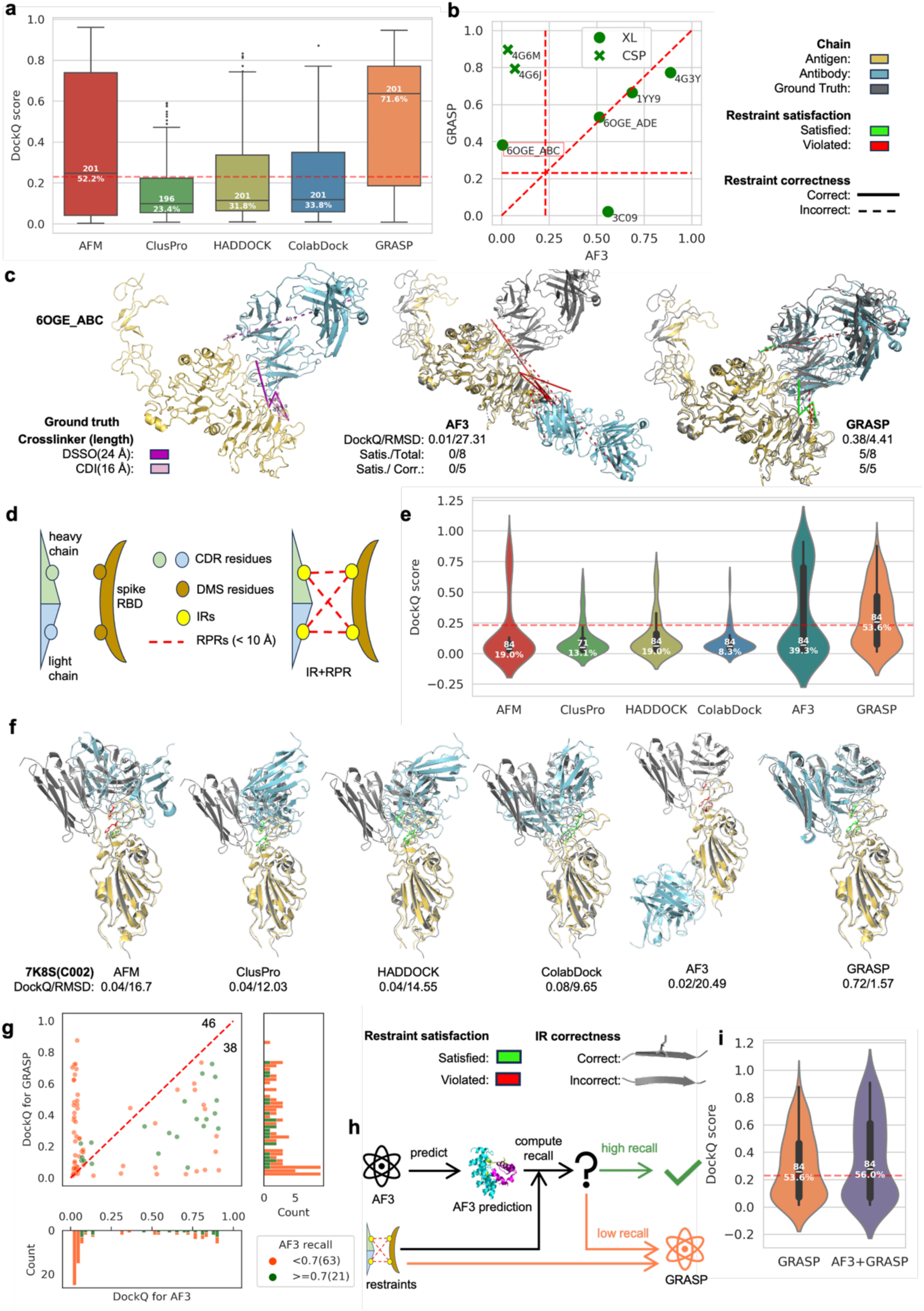
GRASP increases antigen-antibody structure prediction accuracy. **a.** Boxplot for DockQ distributions of various methods across all antigen-antibody complexes in BM5.5. The central line in each box indicates the median, the box edges show the quartiles, and the whiskers extend to 1.5 times the interquartile range, with points beyond this being outliers. The numbers above the medians represent the number of cases. ClusPro fails in some cases, leading to fewer case number. The numbers below the median lines indicate the success rate, in which cases with DockQ ≥ 0.23 are considered successful. The red dashed horizontal line indicates DockQ = 0.23 (same for subsequent panels). **b.** Scatter plot comparing the prediction accuracy of GRASP and AF3 for antigen-antibody complexes with a single type of experimental restraint. The slanted, horizontal, and vertical red dashed lines represent *y* = *x*, *x* = 0.23, and *y* = 0.23 respectively. The case highlighted within the yellow box is shown in detail in panel (**c**). **c.** 3D visualization of the GRASP-predicted structure and AF3-predicted structure for 6OGE_ABC. The left subfigure shows the two types of crosslinkers in the ground truth. The middle and right ones display the AF3 and GRASP predicted structures. The predicted structures are colored cyan (antibody) and yellow (antigen), with the latter superimposed on the real structure in gray (same for subfigure **e**). Satisfied restraints are shown in green and violated ones in red. Correct RPRs are connected by solid lines, while incorrect RPRs are shown with dashed lines. **d.** Schematic representation of the restraint strategy used by GRASP on the experimental COVID-19 DMS dataset. **e.** Violin plots of DockQ scores for various methods on the experimental DMS dataset. **f.** 3D structure visualization of various methods on 7K8S (antibody: C002). For simplicity, only the DMS sites are shown, with correct predictions displayed as sticks and incorrect ones as cartoons. Satisfied restraints are shown in green, and violated restraints in red. **g**. Scatter plot comparing DockQ scores of AF3 and GRASP on samples with high (orange) and low (green) AF3 recall (DMS restraint satisfaction rate). The red dashed line represents *y* = *x*, with the numbers on either side of the line indicating the number of samples falling on each side. The numbers in parentheses in the legend represent the total number of samples for each category. **h.** Schematic illustration of the strategy combining AF3 and GRASP based on DMS restraint satisfaction rates. **i.** Violin plot of DockQ score distributions for the combined strategy (AF3+GRASP) versus GRASP alone.

We further evaluated the performance of GRASP assisted by experimental data. This includes 7 CSP or XL samples from the previous section and 84 additional DMS samples. AF3 are performed on all these samples. The results (Fig. 4b, Table S3) show that, GRASP outperformed AF3, achieving a success rate of 6/7 compared to AF3’s 4/7. For the only case where GRASP failed with PDB ID of 3C09 (Fig. S9), 10 out of 12 experimental restraints were not satisfied by the X-ray structure, suggesting the possibility that this complex has an alternative binding conformation. Specifically, in the case of 6OGE_ABC in which three out of eight XL restraints are incorrect (Fig. 4c), GRASP outperformed AF3. The DockQ scores are 0.38 and 0.01, respectively. GRASP’s prediction met all correct restraints and correctly ignored the incorrect ones, again demonstrating its resistence to experimental noise.

To validate GRASP’s performance on DMS data, we additionally compiled a new DMS dataset from experimental collection^41^, which consists of 84 different monoclonal antibodies bound to the Receptor Binding Domain (RBD) of the SARS-CoV-2 spike protein (see Methods). We selected residues with escape scores above 0.2 as the antigen-side interface to reduce noise (Fig. S10). We used complementarity-determining region (CDR) residues as the antibody-side interface and applied residue pair restraints across the interface residues on both sides (Fig. 4d). GRASP outperformed all other methods with a DockQ median of 0.25 and a success rate of 53.6%. In comparison, for AF3 the DockQ median is 0.07 and the success rate is 39.3% (Fig. 4e, Table S5).

For 7K8S (Fig. 4f), mutations at three sites in the RBD led to immune escape from the monoclonal antibody C002. AFM and AF3 performed poorly, and AF3 misidentified the binding site entirely. ClusPro, HADDOCK, and ColabDock met DMS restraints but failed to predict the correct binding orientation. GRASP, in contrast, corrected the interface in assistance of DMS data and accurately predicted the structure with a DockQ score of 0.72.

It is notable that AF3 excelled in some cases especially for those with DockQ>0.23 (Fig. 4g), and outperforms GRASP in 38 out of 84 cases. Among those 16 cases have over 70% agreement with DMS restraints. To take advantage of both GRASP and AF3, we proposed a joint protocol (Fig. 4h): Use AF3 for initial predictions and accept structures that align well with experimental restraints, otherwise use GRASP to improve predictions by restraints. This approach improved the overall prediction to a median DockQ score of 0.28 and a success rate of 56.0%, surpassing both GRASP (0.25; 53.6%) and AF3 (0.06; 39.3%) (Figs. 4e,i).

### GRASP can serve as a toolkit for integrative modeling

Single experimental technique might provide limited structure information, making it difficult to determine protein interaction completely. Integrating multi-source experimental restraints to assist modeling has become a promising way to study the structure and function of protein complexes^42^. Since GRASP simultaneously supports the input of multiple types of restraints, it can naturally serve as a tool for integrative modeling. We further expand the applicable restraint types of GRASP to include cryo-EM density map by iteratively docking the GRASP-predicted structure into the density map with our GPU-accelerated program Combfit and then extract the spatial arrangement of subunits as contact restraints to guide the GRASP prediction. The protocol is illustrated in Fig. S11. Since high resolution density map is usually sufficient enough to resolve atomic structures with specialized de-novo atomic model building softwares like ModelAngelo^43^, we here only integrate low-to-medium resolution maps. We collected several cases with different combinations of varied experimental approaches to demonstrated the usage and performance of GRASP for integrative modeling.

The first case contains experimental information obtained by XL, mutagenesis, and density map. It characterizes human APOBEC3G (A3G) proteins bound to the substrate receptor module of CRL5 containing HIV-1 Vif, CBFβ, Elongin B and Elongin C (VCBC). We first applied the mutagenesis data between A3G and Vif and XL-MS data containing 13 pairs of cross-linked peptides^44^ to model this system. To validate the prediction, we searched against PDB database and identified a ground truth complex structure released after Robyn et al^44^ (see Methods). 11 of 13 crosslinked residue pairs satisfy the maximum *C*_α_ **-***C*_α_ distance of 30 Å in the ground truth structure^45^ (Fig.5a, left panel). AFM mispredicted the binding interface between A3G and Vif with four pairs of unsatisfied XL restraints. When integrating XL and mutagenesis data, GRASP-predicted a structure satisfying all interface restraints and 12 out of 13 XL restraints with a mean DockQ of 0.435, but A3G also took a wrong orientation. This is partly due to the ambiguity of the restraints. The two K2/A3G-K158/Vif and K2/A3G-K160/Vif restraints biased GRASP to pull A3G closer to the loop region(155-160) in Vif. This case study shows that although GRASP has a good ability of integrating restraints information, it can also be misled by restraints of low quality.

We then added cryo-EM density map from Li et al^45^ to evaluate whether additional information can improve the model performance. The original cryo-EM density map from which the groud truth structure was resolved has a resolution of 2.7 Å. We applied low-pass guassian filter function in ChimeraX^46^ with a width of 5 Å to the density map to obtain a lower resolution of 7.82 Å measured by EMAN2^47^ and integrated the medium-resolution density map information using our protocol. After utilizing the additional density map information, GRASP accurately predicted the binding interface between Vif and A3G, achieving a TM-score of 0.888 and an overall mean DockQ score of 0.445. This predicted structure also maintains a high restraint satisfaction rate (Fig. 5a, right panel).

**Fig. 5.**
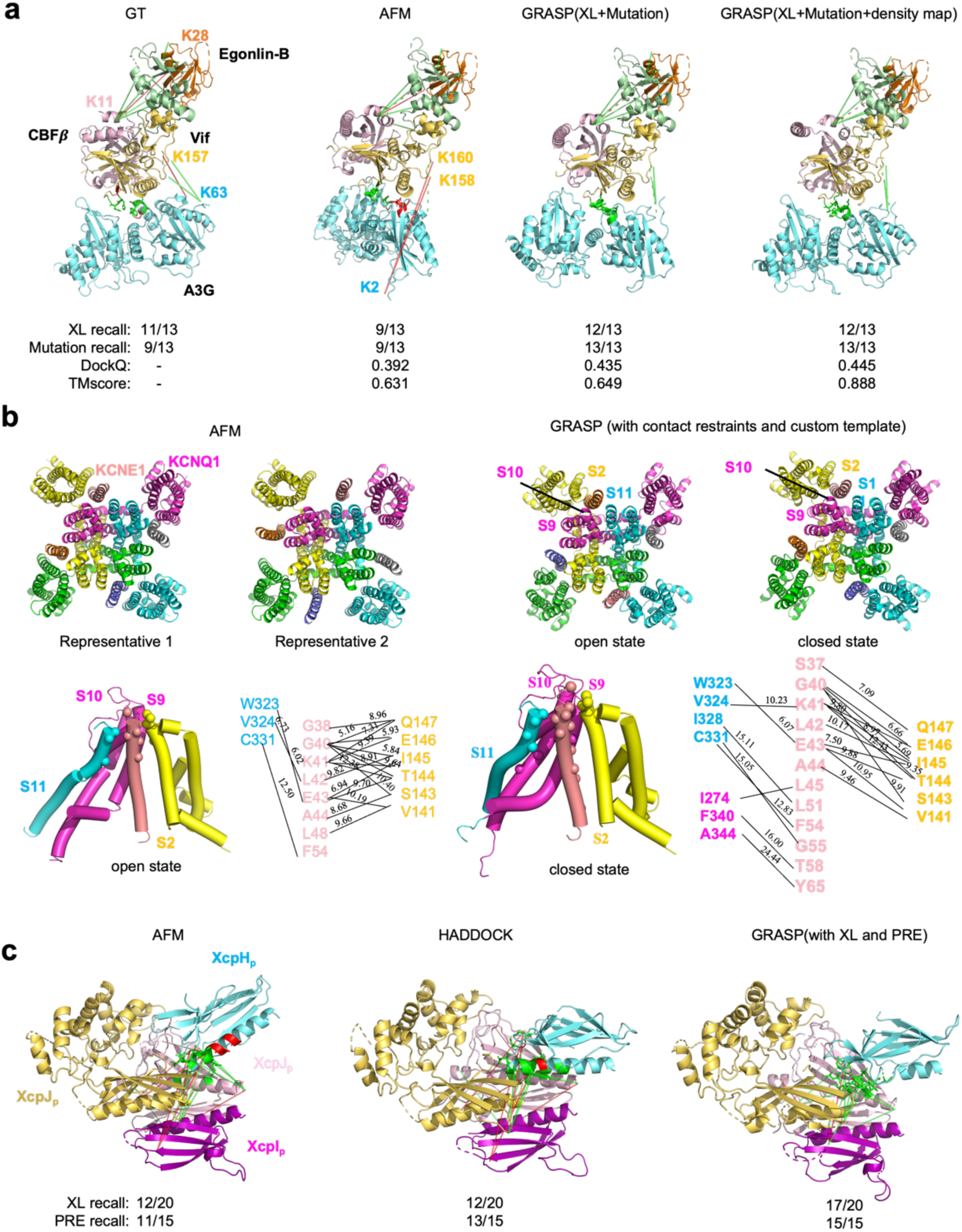
Applying GRASP in integrative modeling results in reasonable structures. **a.**Visualization of experimental restraints for A3G-VCBC complex on ground truth(pdb: 8CX0, left), AFM predicted structure (middle left), GRASP-predicted structure using XL and mutagenesis (middle right), and GRASP-predicted structure using XL, mutagenesis and density map(right). Green lines refer to satisfied XL restraints and red lines refer to violated ones. Dash lines refer to XL restraints which are violated in ground truth in panels **a** and **c**. In this panel, XL recall refers to the ratio of restrained residues pairs with *C*_α_ -*C*_α_ distance below 30 Å. Mutagensis recall refers to ratio of restrained residues whose *C*_"_ atoms are within 10 Å of the *C*_"_ atoms of residues from another chain. **b.** Predicted structures for KCNE1-KCNQ1 complex from AFM (upper left, with two representative structures drawn) and GRASP using the contact restraints and custom templates (upper right). S1-S11 refer to the helices arranged from N terminal to C terminal. The down panels are enlarged interface between KCNE1 and KCNQ1 predicted by GRASP for the open (down left) and closed (down right) state. *C*_α_ -*C*_α_ distance computed for each the utilized RPRs are labeled. Two violated restraints are K63/A3G-K157/Vif and K11/CBFβ-K28/ElonginB. **c**. Visualization of modeling structure of XcpHIJK from AFM, HADDOCK and GRASP using XL and PRE data. XL recall refers to the ratio of crosslinked residues pair with *C*_α_ -*C*_α_ distance below 30 Å. PRE recall refers to the ratio of restrained residues whose *C*_"_ atoms are within 10 Å of the *C*_"_ atoms of residues from another chain.

In the second case, we utilized experimental information from disulfide crosslinking, Cd(II)-cysteine bridging and double mutant cycle analysis from Georg et al^48^ to model the open and close state of voltage-gated potassium(K) channel KCNQ1 and its regulatory protein KCNE1 with stoichiometry 4:4. These restraints were organized in a state-dependent manner and unified into RPRs. The structures of KCNQ1 from KCNQ1-KCNE3 complex^49^ in open or closed state were used as single chain templates to bias GRASP’s prediction toward the corresponding state. Notably, AFM predicted structures are similar to the closed state with low confidence on the position of KCNE1. Different random seeds produced an ensemble of KCNE1 binding across the whole surface of KCNQ1 with two representative interfaces shown in Fig. 5b (left two panels). By integrating multi-source RPRs and customed single-chain templates, GRASP successfully predicted the binding interface in both states. The GRASP-generated structures satisfied the majority of the restraints in both states except two restraints in the closed state: F334/KCNQ1-T58/KCNE1 and A344/KCNQ1-Y65/KCNE1 (Fig. 5b, down panel). The predicted conformation of the open and closed state showed minimal differences. Compared with the open state, the S11 tail of KCNQ1 twisted more towards the center of the ion channel in the closed state, and there are more contacts between KCNE1 and the S9 and S10 helices of KCNQ1. Moreover, the tail of KCNE1 twisted more towards the S2 helix of KCNQ1 in the closed conformation.

The available experimental data for the third case contains a set of XL data and NMR Paramagnetic Relaxation Enhancement (PRE) data from Escobar et al^50^. We used this data to model the type II secretion system, where XcpH is assembled into the trimeric initiating tip complex XcpIJK. Since PRE experiments were conducted between XcpH and XcpJ, both XL data and PRE data were converted into RPRs. The AFM-predicted structure satisfies 15 out of 20 XL restraints and 11 out of 15 PRE restraints with one end of the N-terminal helix of XcpH positioned far from XcpIJK (Fig. 5c, left panel). Esobar et al^50^ built a model using HADDOCK where the helix of XcpH is arranged parallel to the XcpJ helix and the structure satisfies 15 out of 20 XL restraints and 13 out of 15 PRE restraints(Fig. 5c, middle panel). GRASP-built structure is similar to the structure built by Esobar et al^50^ but the helix of XcpH is slightly apart from XcpIJK, which satisfies more restraints with 18 out of 20 for XL and 15 out of 15 for PRE (Fig. 5c, right panel).

### GRASP provides structure insight for dynamic protein-interaction network in mitochondria

Traditional structure determination methods such as X-ray and cryo-EM require purified protein complexes and stable binding between protein subunits. Transient and dynamic protein-protein interactions are hard to capture. Recently, chemical crosslinking technique was improved to detect protein interactome even in whole-cell scale^33,51^. The captured interactions by in-cell XL-MS provide important information of *in situ* protein-protein interaction but lack the details. Here, we utilized GRASP to build the structure models for these interactions.

In the following example, we collected XL-MS data of Chen et al^35^ who developed a targeted delivery system to in-situ map protein interactions in mitochondria. The 158 inter-protein crosslinks were mapped to 144 protein interactions using human MitoCarta 3.0 database^52^. We transformed these cross-linked peptides into RPRs with distance upper limit of 30 Å to predict protein interactions in-batch. Protein interactions with predicted structures fail to satisfy restraints or with pLDDT below 75 were filterd out for quality control. Notably, only three PPIs were discarded due to unsatisfied restraints, which are either with low string score (0.209 for PHB2-MCAT) or unannotated (for LETM1-TRAP1 and TUFM-ACACB). After filtering we obtained 119 2-body protein interactions with 62 co-located in the same subcellular localization (Fig. 6a). 3, 22, and 37 PPIs located in the mitochondrial outer membrane, inner membrane, and matrix, respectively. It can be observed that PPIs in the mitochondria matrix tend to be gobular and are enriched in short helices while those in the membrane have long helix and beta barrel since they are embeded in lipid bilayer. The remaining 57 pairs of PPIs which are involved in cross-compartment interactions of mitochondrial and their interactions are likely highly dynamic and transient (Fig. S12).

**Fig. 6.**
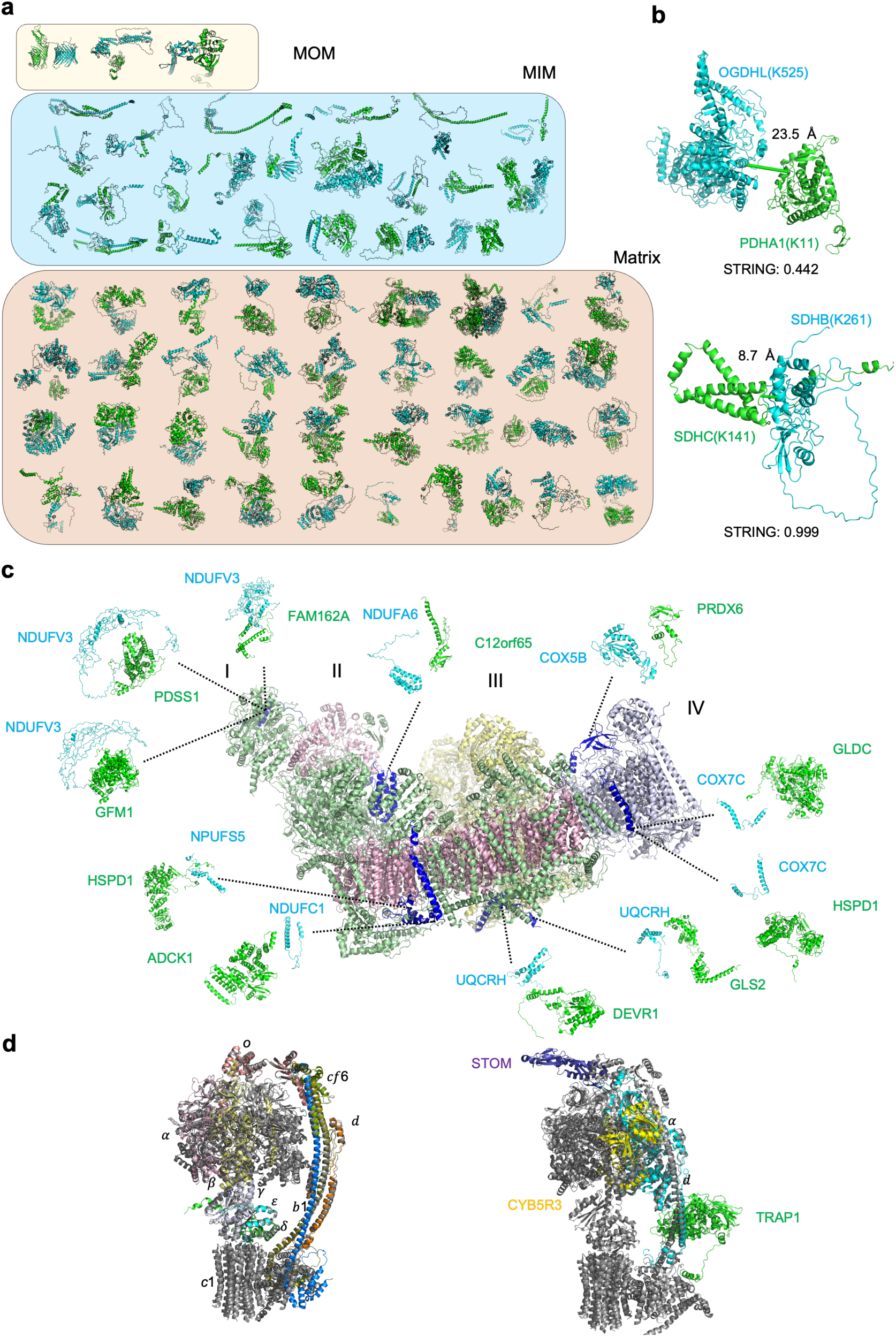
Dimeric structure models are predicted by GRASP for PPIs captured by targets XL-MS in mitochondria. **a.** Visualization for PPIs co-located in the same subcellular localization for mitochondria outer membrane (upper), inner membrane (middle) and matrix (lower). **b.** Two PPIs (OGDHL-PDHA1, upper and SDHB-SDHC, lower) in the tricarboxylic acid cycle. Green lines refer to the well-satisfied restraints and red lines refer to violated restraints. **c.** Visualization of potentially dynamic PPIs between OXPHOS complex I-IV and other proteins. Complex I-IV were colored in palegreen, lightpink, paleyellow and lightwhite,respectively. The crosslinked chain in the Complex I-IV were colored in deep blue. Chains of predicted PPIs were colored in cyan and green **d.** Visualization of PPIs related to F0-F1 ATP synthetase. PPIs within ATP synthetase are shown on the left panel and representative PPIs between ATP synthetase and other proteins are shown on the right panel. The ground truth structure of ATP synthetase (PDB:8H9V) was colored in grey and corsslinked chains in ATP systhease were in different colors.

The targeted XL-MS data captured 2 pairs of PPIs enriched in the TCA (tricarboxylic acid cycle) cycle. One of them is K525 of 2-oxoglutarate dehydrogenase-like (OGDHL) linked to K11 of pyruvate dehydrogenase E1 component subunit α (PDHA1), with a *C*_α_ **-** *C*_α_ distance of 21.5 Å in the GRASP-predicted structure (Fig. 6b). GRASP predicted a relatively weak interaction between OGDHL and PDHA1 which is supported by the confidence score of 0.442 in the STRING database^53^. The second one connects K141 of the succinate dehydrogenase cytochrome b560 subunit (SDHC) with K261 of the succinate dehydrogenase [ubiquinone] iron-sulfur subunit (SDHB), with a *C*_α_ **-***C*_α_ distance of 8.7 Å in the GRASP-predicted structure. SDHC wraps around SDHB and forms a stable interaction with a STRING^53^ confidence score of 0.999.

Oxidative phosphorylation (OXPHOS) is mediated by enzyme complexes, known as complexe I to V, which are responsible for producing most of the ATP required for normal cell function. Most of the captured interactions between complex I-IV and other proteins participate in the process of assembling OXPHOS system, mitochondrial protein synthesis and fatty acid metabolism accoding to the annotation of NCBI^54^. We show the interaction between complex I-IV and other proteins in Fig. 6c. NDUFV3 is a protein from Complex I and with 3 peptides connecting it with other proteins. Interestingly, NDUFV3 adopts a different conformation upon interaction with FAM162A, a protein active in the process of apoptosis^55^, compared to the predicted structures of the other two NDUFV3-related PPIs.

F0-F1 ATP synthetase (complex V) is the most cross-linked complex possessing 11 restraints within the complex and additional 11 restraints between ATP synthetase and other proteins. Although we only modeled two-body interactions, 6 out of 7 PPIs align well with the reference structure across *α*, *β*, *γ*, *δ*, *ε*, *o* subunits in F1 and *b*1*,d*,*cf*6 subunits in the rod (Fig. 6d). In addition, we built three interfaces between other proteins and the mature F0-F1 ATP synthase. TRAP1 binds to *d* subunit while STOM and CB5R3 bind to *α* subunit at different sites. The relative orientation between TRAP1 and *d* subunit is consistent with the location of TRAP1 in the matrix. Similarly, for STOM and CB5R3 located in the inner membrane, their predicted orientations relative to ATP synthetase also align with their location. TRAP1 is believed to function by balancing the oxidative phosphorylation and aerobic glycolysis^56^ while STOM may influence the activity of ion channel according to GO annotation^57^ and CYB5R3 serves as NADH-cytochrome b5 reductase^58^. These predicted structures may further assist discovery of the cause of diseases when resolved structures are absent.

## Discussion

In this work we propose GRASP, a generalized model that integrates a variety of experimental restraints to enhance protein complex structure prediction. By incorporating RPR and IR information directly into the GRASP model and fintuning both the AFM and the self-trained MEGAFold-Multimer checkpoints with additional restraint-related loss terms, GRASP can simultaneously process one or multiple types of experimental data as restraints to improve the accuracy and reliability of predicted protein complex structure. This approach allows the integration of various data sources to complement and/or cross-validate each other. An iterative noise filtering process is also introduced to reduce the impact of errorous restraints. GRASP uses a model inference approach and only needs to be trained once to obtain a minimal inference time, offering the advantage of supporting diverse restraint forms especially for large complexes. Compared to other docking methods, GRASP does not require monomer or complex structures as templates.

GRASP supports both residue pair restraints at various distances and interface restraints. It demonstrates excellent performance in complex structure prediction by effectively integrating various simulated and experimental restraint data. Under the simulated XL or real-world XL, CL, CSP restraints, GRASP achieves higher prediction accuracy than other methods, demonstrating its efficiency in integrating independent data types. For the crucial but highly challenging antigen-antibody prediction task, GRASP leverages sparse restraints from CSP, XL, or DMS data to surpass AF3 in accuracy. Additionally, we propose a combined strategy with AF3 to further enhance prediction performance.

GRASP naturally serves as a tool for integrative modeling. By combining the varied information across cryo-EM, XL, NMR PRE, mutagenesis, and Cd(II)-cysteine bridging experiments in three cases, we demonstrated the potential for GRASP to perform the complicated restrained modeling tasks. Further, with this tool, we are able to model structures with actively developing *in situ* or dynamic detection techiques including in-cell XL, solution NMR, *in situ* cryo-ET, helping reveal molecular mechanisms near physiological conditions.

In the post-AlphaFold stage, structural biologists are focusing on more complex protein interactions with essential biological applications. GRASP can act as a new toolkit for integrating experimental information with different levels of structural details and accuracy. We are optimizing the integrative strategy of GRASP to enable solution of more challenging application scenarios like transcription complex, ribosome, spiceosome and various membrane protein complexes. Moreover, GRASP can also be consolidated into the workflow of existing integrative modeling tools like CombFold^59^ thus combining the advantages of sampling/searching-based methods and deep learning-based methods.

## Method

### Datasets

#### Training dataset

We used the PSP dataset^60^ as the basis for the GRASP and MEGAFold-Multimer training set. The PSP dataset contains a 570k true structure set released before October 8, 2021 with protein sequences, experimental structures, MSAs, and templates, along with a 760k distillation set with predicted structures, MSAs, and templates. Complex samples were constructed by pairing single chain MSAs with the same PDB id in the true structure set using the pairing protocol within AFM^4^ and the rest single-chain information combined.

#### Self-curated test dataset

To assess the ability of GRASP to integrate two types of restraints (contact-RPR and IR), we curated a test dataset and benchmarked GRASP against several other methods.

##### Sample collection

To ensure a comprehensive and unbiased evaluation, we created a new challenging and high quality dataset to avoid data leakage and redundancy. We retrieved the protein sequences for each PDB ID released before March 20, 2023, split them into individual chains, and excluded non-protein chains and those shorter than 20 residues, resulting in about 707,901 single-chain protein sequences. Using “easy cluster” in MMSeqs2^61^ with a 70% sequence similarity threshold, we clustered these sequences into 58,671 clusters.

To avoid data leakage, we removed all protein complexes released before October 8, 2021, as well as complexes with the same cluster composition as those released before this date. We note here that we considered only the cluster type; for example, if a complex with 3 chains from cluster A was released before October 8, 2021, a complex with 2 chains from the same cluster would be excluded. Only the biological assembly 1 was used. We then applied the following criteria to retain complexes: 1) total chain length ≤ 5120 residues; 2) resolution better than 3.0 Å; 3) 2 to 20 chains; 4) shortest chain ≥ 32 residues; 5) every chain in contact with at least one other chains from the complex; 6) for duplicates with the same cluster composition, the complex with the smallest PDB ID was kept. This process resulted in 1,336 protein complexes. We further refined this set to 313 complexes where the AFM prediction had a DockQ score below 0.23.

We focus solely on interfaces in this part. For complexes with three or more chains, we randomly sampled up to five non-redundant pairs of contacting chains, resulting in a total of 713 interfaces.

##### Contact-RPR restraint sampling

The 313 protein complexes were randomly divided into three groups, each interface in the three groups was assigned with 2, 5, or 10 contact-RPRs, respectively. Contact-RPRs were sampled from inter-chain residue pairs with *C*_*β*_ distances within 8 Å, and no erroneous restraints were introduced.

##### IR sampling

Similarly, the 313 protein complexes were randomly divided into three groups, with interfaces in each group assigned with 4, 10, or 20 IRs. IRs were randomly sampled from interface residues within 8 Å of the other chain, without introducing erroneous restraints.

#### Simulated XL dataset

To assess the performance of GRASP on simulated XL data, we generated crosslinks (XLs) for the 313 complexes curated in the previous section. We performed inter-chain XL simulations using Xwalk^62^, applying a *C*_*β*_ -*C*_*β*_ cutoff of 25 Å and trypsin digestion. Two sets of restraints were simulated: one with XLs amounting to 2% of the total sequence length, and the other with 5%. To mimic real experimental conditions, we included 20% erroneous restraints (FDR = 20%), defined as residue pairs with *C*_*β*_ -*C*_*β*_ distances exceeding 25 Å, where at least one residue is Lys, Ser, Thr, or Tyr. After excluding 19 complexes having fewer than 40% of the desired correct XLs, 294 complexes remained for analysis.

#### Experimental datasets with single restraint type

We constructed datasets with single experimental restraint types to compare the performance of different methods. These include 5 CL, 7 XL and 4 CSP cases.

##### XL dataset

The dataset consists of 5 antigen-antibody complexes^34,35^ (1YY9, 3C09, 4G3Y, 6OGE_ABC and 6OGE_ADE) and 2 large protein complexes^36,37^(4B2T and 5L4K). XLs were transformed into RPRs with distance range determined by the specific crosslinking reagents, e.g. 30 Å for DSSO (see Table S6 for the XL distances for different reagents). For 6OGE, We predicted HER2 extracellualr domain binding to pertuzumab and trastuzum respectively, and named the two complexs as 6OGE_ABC and 6OGE_ADE following Wang et al^34^. 4B2T has a full length of 8682 residues which is too large for our model. Since it is of C2 symmetry, we chose to make structure prediction on the non-symmetric half.

##### CL dataset

The 5 cases of covalent-labeling are collected from Drake^23^(1YAG, 2F8O, 4INS4, 4INS8 and 4INS12), as did in our ColabDock model^22^. Covalent labeling data provides information on residues buried in the binding interface and can be naturally converted into IRs. The prediction results of ColabDock, HADDOCK and ClusPro are available from Ref^22^.

##### CSP dataset

We obtained four CSP examples, two from Dominguez^25^ (1GGR and 3EZA) and the other two from Blech^38^ (4G6M and 4G6J). The chemical shift perturbation data are transferred into the form of IRs. Since 4G6M and 4G6J are antibody-antigen systems and only have restraints on the antigen side, we selected the residues in the CDR region as IRs on antibody side. The antibody CDR regions were identified using the Chothia scheme via http://www.bioinf.org.uk/abs/abnum/^63^. The prediction result of 1GGR and 3EZA of ColabDock, HADDOCK and ClusPro are acquired from Feng et al^22^. For 4G6M and 4G6J, We ran calculations using these tools with the same settings as used by Feng et al^22^ to complete the benchmark.

#### Simulated DMS dataset

To evaluate GRASP on antibody-antigen systems, we simulated three sets of DMS restraints for each of the 67 antibody-antigen complexes in the BM5.5^40^ dataset. Of these, 45 complexes had already been assessed by Feng et al^22^, and we reused the simulated restraints for them. The remaining 22 complexes were processed using the same restraint simulation strategy as Feng et al^22^.

#### Experimental DMS dataset

We obtained deep mutational scanning profiles for 3051 SARS-CoV-2 WT RBD-targeting monoclonal antibodies (mAbs) from Cao et al^41^. We then searched the PDB^3^ database using the mAb names to identify protein complexes that met our criteria, specifically those containing the spike glycoprotein, light chain, and heavy chain. This search resulted in a final set of 84 complexes, with additional details provided in Table S5. The spike glycoproteins were aligned to the Wuhan-Hu-1 RBD sequence (residues N331 to T531, GenBank: MN908947), and mutation escape sites for each complex were determined. We trimmed the spike protein to include only the RBD regions for further analysis. The antibody CDR regions were identified using the Chothia scheme via http://www.bioinf.org.uk/abs/abnum/^63^.

#### Integrative modeling dataset

For A3G-VCBC complex, we collected XL and mutagensis data from Robyn^44^ and only kept the chains which are present in the ground truth. We additionally removed intra-chain XL restraints and those involving residue types that cannot be cross-linked by the crosslinking reagent. The remaining XLs were transformed into RPRs with distance upper limit of 30 Å according to the reagent DSSO. Mutagensis data were transformed into IRs. Finally, we obtained a sample with 13 pairs of XLs and 13 mutagenesis IRs. We used guassian filter function in ChimeraX with width 5 Å to raw map with resolution 2.7 Å and obtained a medium-resolution map to assist prediction. In addition, we separated the complex into two parts of A3G and VCBC complex to run Combfit.

For the KCNE1-KCNQ1 complex, we curated the experimetal restraints from Kuenze et al^48^ including disulfide crosslinking, Cd(II)-cysteine bridging and double mutant cycle analysis in either state to model. These restraints were transformed to RPRs and the expected distance range following the setting of Kuenze et al^48^. This procedure led to 19 RPRs for open state and 22 RPRs for closed state. These restraints were assigned to all KCNE1-KCNQ1 combinations. Moreover, we used the the single chain of KCNQ1 from PDB entry 6V01 and 6V00 as the template to assist prediction for open state and closed state, respectively.

For XcpH-IJK complex, we obtained acidic XL and PRE data from Ref^50^. These restraints were both transformed into RPRs of different distance ranges. We set maximum *C*_*α*_ -*C*_*α*_ distance to 25 Å for acidic XL and 10 Å for the PRE restraints.

We developed a pipeline to retrieve experimentally resolved structures for the above three complexes. We first searched for structures corresponding to the the query complex sequence using the “rcsbsearchapi”, a Python interface for the RCSB PDB search API^64^. By performing protein sequence search, we identified all PDB entities that shared at least 85% sequence identity with the target complex, applying an E-value cutoff of 1e-3. The restraint recall for each entity was then calculated based on experimental restraints, and entities with recall below 0.75 were filtered out. The structure containing the most consistent chains with the modeling system was selected as the ground truth structure. If multiple candidate structures were identified, the one with the highest resolution was selected. By doing so, we identified the 8CX0 as the ground truth for A3G-VCBC complex and failed to find suitable ground truth for other two complexes.

#### Mitchondria dataset

We collected targeted XL-MS data from Chen et al^65^ and kept only the inter-protein XL restraints. We merged XL restraints with the same gene name to improve the restraint quantity for each pair of PPIs. We renumbered the crosslinked sites according to corresponding sequence in human MitoCarta 3.0 database^52^ by sequence alignment and prepared the sequences for structure prediction. This led to a dataset including 144 PPIs with 1-3 pair RPRs for each. We set maximum *C*_*α*_ -*C*_*α*_ distance to 30 Å for DSS to be consistent with Ref^65^.

### Model training strategy

#### MEGAFold-Multimer training

We trained a MEGAFold-Multimer model from scratch using 128 Ascend-910A (32G) AI chips. MEGAFold-Multimer is a reimplementation of AFM within the MindSpore framework. The training process consisted of two stages (46,200 total steps over 17 days): an initial training phase (33,000 steps over 12 days) and a fine-tuning phase (13,200 steps over 5 days).

In the initial training phase, the learning rate started at 0 and increased to 3e-4 with a 1,000-step warmup, then decayed to 3e-5 using cosine decay over 74,000 steps and stopped at 33,000 steps. During fine-tuning, the learning rate began at 5e-5 and decayed to 5e-6 using cosine decay over 75,000 steps, stopping at 13,200 steps. The sampling ratio from the distillation dataset, true structure dataset, and multimer dataset was 5:2:3 during the initial training stage and 3:2:5 during the fine-tuning stage. The crop size was set to 384. All other configurations followed those of AFM.

We evaluated MEGAFold-Multimer on 172 non-redundant complexes released between July 20, 2022 and March 1, 2023, demonstrating a comparable accuracy to AFM (Fig. S1).

#### Fine-tuning the GRASP model

We fine-tuned on both AFM v2.3 Model-1 checkpoint and MEGAFold-Multimer checkpoint over 9 days separately using 64 Ascend-910A (32G) AI chips. To achieve a more stable prediction, we generated five models. Four of these were fine-tuned from AFM v2.3 Model 1, taken at 8k (JAX-8k), 14k (JAX-14k), 20k (JAX-20k), and 22k (JAX-22k) steps during the fine-tuning process. The fifth model (MS-22k) was fine-tuned 22k step from the MEGAFold-Multimer model.

We adjusted the multimer/monomer ratio to 4:1 in GRASP training, set the learning rate to 1e-4, and used a crop size of 384. All other configurations were consistent with those of AFM unless specified otherwise.

#### Restraints simulation during fine-tuning

##### Restraint number

We simulated intra-chain RPRs (intra-RPRs), inter-chain RPRs (inter-RPRs), and IRs during fine-tuning, separately. Intra-RPR and inter-RPR were handled separately considering their differences in distance distribution, quantity, and impact on the predicte structure. First, we randomly and independently sampled the numbers (*n*) of the three types of restraints for each case. For each restraint type, there is a 50% probability that *n* equals 0, and a 50% probability that *n* is sampled from a piecewise function normalized across the restraint number *n*. The piecewise function is defined as follows:

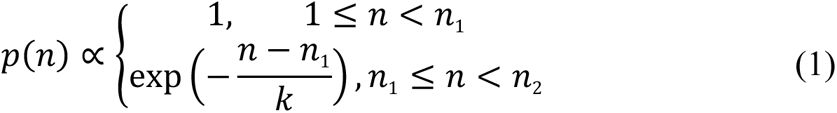

From 1 to *n*_1_ , *n* is uniformly distributed; from *n*_1_ to *n*_2_ , an exponential decay function is applied, with the decay rate controlled by parameter *k*. In GRASP finetuning, the values of *n*_1_ /*n*_2_ /*k* are set as following for each restraint type: 40/80/8 for IR, 20/40/4 for inter-RPR, and 80/160/16 for intra-RPR. If the actual number of restraints is less than the sampled number *n*, the final count is set as the actual number.

##### Sampling order

To prevent interference between inter-RPR and IR, we carefully designed the sampling process (Fig. S13). We first sampled intra-RPR to a specific number from residue pairs with sequence index separation of 6 or higher. Next, we randomly selected either inter-RPR or IR with equal probability, and then sampled the remaining type, excluding previously selected residues. In each step, sampling was conducted without replacement until the target number of restraints was reached or all candidates were exhausted.

##### RPR restraints

To encode distance information for RPR, we discretized the distance space into a distogram of 30 bins. The 28 central bins cover distances from 4 Å to 32 Å, with each bin spanning 1 Å, while the first and last bins represent distances smaller or larger than distances in this range. We generated two main forms of RPRs with the uniformly distributed and sharp distogram (see following section) with equal probability during training, incorporating 5% erroneous restraints (FDR=0.05).

##### Uniformly Distributed Distogram

This distogram applies a uniform distribution below a variable cutoff randomly selected between 8 and 30 Å for each complex, mimicking the effects of crosslinkers with varying lengths. The sum of the bins below the cutoff equals 1−FDR, while the sum above the cutoff equals FDR.

##### Sharp Distogram

This distogram restrains the residue pair distance to a single bin, where the value for that bin is 1−FDR and the remaining bins sum to FDR. This approach helps the model to learn the specific relationship between distances and their respective bins. We randomly sampled distances to ensure a roughly equal representation across all bins.

##### IR restraints

Interface restraints are binary, where a value of 1 indicates that a residue is at the interface, meaning at least one residue from another chain is within 8 Å. A value of 0 signifies a lack of experimental data on its involvement in the interface. To ensure balanced representation of interface residues on each chain, we sampled these restraints with probabilities inversely proportional to the number of interfaces on that chain, without introducing any erroneous restraints (FDR=0).

#### Restraint-related loss terms

We introduced four restraint-related loss terms, including RPR FAPE loss, RPR dRMSD loss, RPR distrogram loss, and the interface loss.

##### RPR FAPE loss

ℒ_*RPR*_*fape*_ measures the Frame Aligned Point Error (FAPE) between restrained residue pairs. For each pair (*i*, *j*) with an experimental restraint, it aligns the backbone local frame of residue i and calculates the coordinate distance error for the jth residue between the predicted structure and ground truth, as shown in Equation (2). The length scale *Z* is 10 Å for intra-chain RPR and 30 Å for inter-chain RPR.

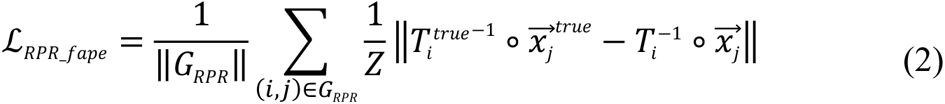

##### RPR dRMSD loss

ℒ_*RPR*_*drmsd*_ quantifies the distance Root-Mean-Square Deviation (dRMSD) associated with restraints. This term computes the RMSD of distances between the key backbone atoms of each restraint residue pair, as shown in Equation (3). Here, *A*_*backbone*_ refers to the four backbone heavy atoms: *C*_*α*_ , *C*_*β*_ (*C*_*α*_ is used here and hereafter for glycine), N and O. The term *d*_*ija*_ represents the distance between atom *a* in residue *i* and the corresponding atom in residue *j*.

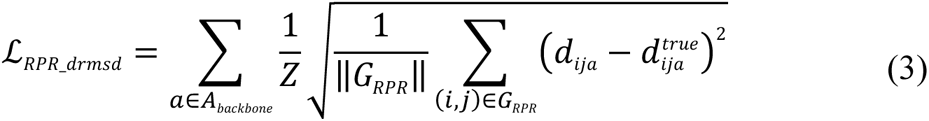

##### RPR distrogram loss

ℒ_*RPR*_*disto*_ encourages the predicted structure to adhere to distogram restraints. Since the distograms show single-peak distributions with a continuous high-probability interval, no penalty is applied if the predicted distance falls within this interval. Otherwise, a penalty is calculated as the clipped minimum distance from the predicted distance to the interval edges with a tolerance *δ*. This term is given in Equation (4), where *a* and *b* represent the interval’s center and length, respectively. The tolerance parameter *δ* was set to 2 Å in our experiment.

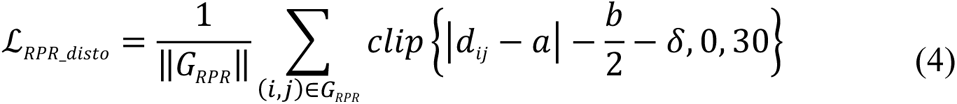

##### Interface loss

For interface loss, we computed the nearest distances from each interface residue *i* to residues in other chains. A clipped linear penalty was applied if this distance exceeded the threshold (8+*δ*) as shown in Equation (5). Here, *I*_*interface*_ represents experimentally observed interface residues, and *C*_*i*_ denotes residues in the same chain as *i*.

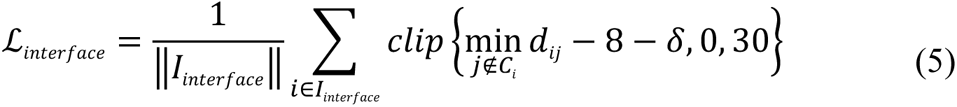

We combined the new loss terms with the original AFM loss. The structure violation loss weight was set to 0.03. Intra-chain and inter-chain RPR losses were computed separately due to their disparity. The final loss function, shown in Equation (6), set the hyperparameters *λ₁*, *λ₂*, *λ₃*, and *λ₄*, to 0.5, 0.05, 0.01, and 0.5, respectively.

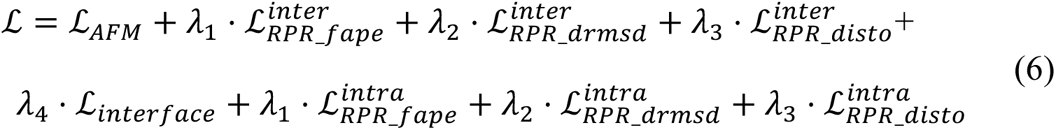

### Iterative retraint filtering

To address noise in experimental restraint signals in model inference, which can lead to inaccurate structure predictions, we implement an iterative restraint filtering strategy. Each round of restraint filtering evaluates the validity of the current restraints based on the predicted structure. Two key metrics are assessed in the order of priority: 1) Chain break at the restrained residues; 2) The extent to which the structure violates the restraint.

We regard the *C*_*α*_ distance between consecutive residues higher than 5 Å as chain break. The neighboring *C*_*α*_ distance (*d*_*NBCA*_ ) for a residue is defined as the average *C*_*α*_ distance to its two (one for terminal residues) flanking residues. The *d*_*NBCA*_ for a RPR restraint is defined as the average *d*_*NBCA*_ of the two restrained residues, while for an IR restraint it is the *d*_*NBCA*_ of the interface residue.

The violation distance (*d*_*viol*_ ) measures how much the predicted structure violates the restraint. For example, if an IR specifies residue i as an interface residue but its closest distance to another chain is 20 Å, *d*_*viol*_ is 12 Å. For an RPR restraint requiring residues i and j to be within 25 Å, *d*_*viol*_ is 0 if their predicted distance is 10 Å, and 5 Å if the distance is 30 Å.

Using these two distances, the irrationality score (*IS*) of a restraint is defined as:

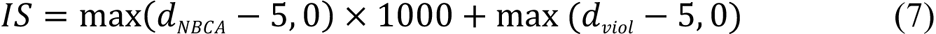

To avoid excessive restraint removal, each iteration filters up to 20% of current restraints, ensuring at least 20% of the original restraints are retained. The filtering stops when the maximum iterations are reached, no restraints are removed, or fewer than 20% of the original restraints remain. IR and RPR filtering are executed independently, and the final predicted structure is then output.

Convergence is typically reached within 2-3 iterations, but a larger number may be needed with a large number of restraints. To improve efficiency, we introduced a "fast inference mode" as shown in Fig. S14. This mode reduces the number of recycles between iterations and recycles the structure after each filtering step, allowing up to 20 inference rounds and making it up to 4 times faster than the standard version. We used this mode in the DMS dataset evaluation, where RPRs across both sides of the interface introduced *m*\**n* RPRs, with *m* and *n* representing the number of interfaces on each side.

### Integrative modeling using Combfit

To smoothly integrate cryo-electron microscopy (cryo-EM) density map infromation into GRASP, we developed Combfit, an advancement over the GPU-based matching software Powerfit^66^, utilizing clustering and genetic algorithms(see details in SI). While ensuring accuracy, Combfit significantly reduces computation time (Fig. S15). Implemented in Python, Combfit offers excellent readability and is easy to be integrated with other models.

In integrative modeling involving cryo-EM, we first converted all restraints, except for the density map, into either RPRs (for XL, and NMR NOESY) or IRs (for CL, mutagenesis, and CSP) to predict the initial structure. The initial structure was then split into single chains or pre-defined chain combinations and docked into density map using Combfit. We next extracted residue pairs from the docked structure where the *C*_*α*_ -*C*_*α*_ distance was ≤25 Å and had decreased by at least 10 Å compared to the initial structure. These residue pairs were ranked by the magnitude of the distance change and the top 500 pairs were selected as candidates. Using a max-min algorithm^67^, up to 30 representive residue pairs were chosen from this set as RPRs. These RPRs, along with the existing RPRs and IRs from other experiments, were then used to guide the GRASP’s second round of prediction. In doing so, we obtained the final structure for intergrative modeling with cryo-EM data.

### Setup for GRASP inference

The AFM protocol^4^ was used to search for MSA and templates and to prepare input features for GRASP. We used reduced databases in cases where HHblits raised errors during MSA searches. The standard iterative retraint filtering strategy was applied with a maximum of 5 iterations for cases with fewer than 3072 residues and 2 iterations for larger ones. Each checkpoint runs with five seeds for complexes under 3072 residues to align with the AFM setup and one seed for larger complexes. For all the predicted structures, we select the predicted structure with the highest pLDDT score as the final prediction, prioritizing those with a recall of at least 30%. Due to memory limitations, the Ascend 910A (32G) could not handle inference for sequences longer than 1280 residues. To address this, we migrated GRASP’s code and checkpoint to JAX (GRASP-JAX). For complexes exceeding 1280 in length, we performed inference using GRASP-JAX on A100 (80G) GPUs. The basic setup described above was used for all GRASP inference, with some unique settings as follows:

For the experimental DMS dataset, we used both RPR and IR (Fig. 4) and applied quick mode iterative filtering to speed up inference (see the “Iterative Restraint Filtering” section). We also evaluated performance when IR and RPR are used separately (Fig. S16).

For the simulated DMS dataset, we divided all complexes into antigen and antibody for docking. We used GRASP in Dimer Mode, which converts complexes with more than two chains into dimers by setting the same “asym_id” for chains in each group and adding a 200-residue gap between chains^59^. This approach improves interface restraint precision, narrows the search space, and simplifies the task. GRASP’s performance without Dimer Mode was also evaluated (Fig. S17). For fair comparison, all other methods were also evaluated by splitting complexes into antigen and antibody.

For the integrative modeling cases, we applied the integrative strategy with Combfit metioned above to incorporate additional information from density map for A3G-VCBC. For the rest two cases standard setup of GRASP was used. For the scoring stage of all three cases, we normalized each structure’s pLDDT and recall by subtracting the mean and dividing by the standard deviation, then selected the one with the highest average normalized score.

For the inference of the dimeric interactions in the mitchondria, we disabled the iterative retraint filtering strategy due to the sparsity of the restraints, the number of which is only 1-3 per complex.

### Configurations for Other Structure Prediction Tools

#### AFM

We used the same input features for AFM as GRASP. Five v2.3 models were used, with each model predicting 5 structures for sequences up to 3072 residues, and one structure for longer sequences due to limitation in resource.

#### AlphaLink

We used the same input features for AlphaLink as GRASP. The AlphaLink-Multimer_SDA_v3.pt checkpoint was used for inference, with the number of random seeds adjusted to generate the same number of structures as GRASP: 25 random seeds for complexes with up to 3072 residues, and 5 seeds for larger complexes.

#### ColabDock

The settings of ColabDock is the same as in Ref^22^. We use AF2^68^ to generate single-chain templates and AFM to generate templates for antibody side. If only one side of the interface was available, the IRs for the other side were generated by selecting residues with solvent accessibility ≥ 40% using the FreeSASA^69^ software. These single-chain templates and processed restraints also serve as inputs of HADDOCK and ClusPro. Please note that since GRASP does not necessarily require single-chain templates and readily handles one-sided interfaces, these two type of information is not needed. For complexes whose total length exceeds 700 residues, we ran segment-based optimization with recommended settings.

#### HADDOCK

We installed a local version HADDOCK 2.4 to perform benchmark experiments. We followed the default setting of HADDOCK and picked the best structure by HADDOCK score from “water” refinement stage as the final structure. For cases which failed in the “it1” or “ water” refinement stage due to energy optimization issues, we picked the best structure in the last stage.

#### ClusPro

We installed the program PIPER provided on the ClusPro website and convert the restraints into json format using offical scripts. We set the restraints satisfication ratio to 50% for interface restraints and decreased this ratio to 30% for cases without available output structure at 50% ratio.

#### AF3

The code for AF3 was not open-source and it is only accessible via the online server (https://golgi.sandbox.google.com/). We used the default settings on the website for our predictions.

### Evaluation metrics

We used four metrics to assess performance: DockQ, TM-score, RMSD, and restraint recall. Residues present in the predictions but missing in the ground truth were excluded in evaluation. For multimers containing homologous chains, we used a greedy strategy in AFM^4^ to match each predicted chain with its best corresponding ground truth chain for all predictions before evaluating.

#### DockQ

DockQ^70^ is a quality measure for docked models based on the CAPRI protocol. Models are categorized as Incorrect (0 < DockQ < 0.23), Acceptable (0.23 ≤ DockQ < 0.49), Medium (0.49 ≤ DockQ < 0.80), and High Quality (DockQ ≥ 0.80). We define cases with a DockQ score above 0.23 as successful, and the success rate is the percentage of such cases in the evaluated dataset. DockQ was calculated for each pair of interacting chains and then averaged for complexes with more than two chains. For antibody-antigen and other complexes already split into dimers in previous studies, such as the five CL real-world cases, we computed DockQ by treating them as dimers.

#### TM-score

TM-score^71^ evaluates structural similarity, focusing on global topology. A score of 1 indicates a perfect match, while scores below 0.17 suggest unrelated proteins. We used the TM-score program (version 2022/02/27) for our calculations.

#### RMSD

RMSD measures the overall structural difference between predicted and reference structures based on *C*_*α*_ atoms.

#### Restraint Recall

Restraint recall measures the percentage of provided restraints satisfied by the predicted structure. RMSD and Restraint recall are computed by custom python scripts.

## Acknowledgement

This work was supported by the National Science and Technology Major Project (No. 2022ZD0115001), the National Natural Science Foundation of China (No. 92053202, 92353304, 21821004 and 21927901) and New Cornerstone Science Foundation (NCI202305). We thank Prof.Yang Kaiguang from Dalian Institute of Chemical Physics to provide guidance for usage of the targeted XL-MS in the mitochondira.

## Data and Code Availability

Data and code of this work will be released soon.

